# Escape from X inactivation varies across genes and tissues and shapes sex-biased sex chromosome gene expression

**DOI:** 10.64898/2025.12.19.695225

**Authors:** Alex R. DeCasien

## Abstract

Sex differences in human health and disease are shaped by complex interactions between hormones, environment, and genetic factors – including those associated with sex chromosomes. While X chromosome inactivation (XCI) in females generally silences one copy of the X to equalize dosage with males, a subset of genes “escape” XCI and remain expressed from both X chromosomes. In this study, I integrate allele-specific expression data from three females with non-mosaic XCI, sex-biased expression profiles from over 40 tissues, and enhancer activity data from GTEx to explore how variation in the magnitude of XCI escape contributes to sex-biased gene expression across the human body. I confirm that female-biased expression on the X chromosome is a poor proxy for escape from XCI. I find that XCI extends into the pseudoautosomal region (PAR) and that the extent of inactivation strongly predicts male-biased expression of PAR genes. Conversely, stronger escape from XCI in non-PAR X-linked (NPX) genes is associated with more pronounced female-biased expression. Across both PAR and NPX genes, escape patterns are shaped by topologically associating domains (TADs) and sex-biased expression is supported by proximity to sex-biased enhancer activity. These findings reveal a direct, tissue-specific relationship between the strength of XCI escape and the magnitude of sex-biased gene expression, providing a mechanistic framework for understanding how the X chromosome contributes to sex-biased biology.

## Introduction

Humans exhibit sex differences in the prevalence and presentation of many conditions. For example, males have higher rates of non-reproductive cancers, early-onset neurodevelopmental disorders, and Parkinson’s disease, while females are more likely to be diagnosed with autoimmune diseases and Alzheimer’s disease [1–5]. These sex differences arise from a complex interplay of hormones, environment, and genetic factors – including the sex chromosomes.

The X chromosome, in particular, plays a central role in shaping sex-biased biology. Genes with sex-biased fitness effects are predicted to accumulate on sex chromosomes over evolutionary time [6–8]. In line with this, X-linked genes exhibit some of the strongest observed sex differences in gene expression [9] and may be enriched for immune and brain functions [10–13] (cf. [14]), two domains that exhibit pronounced sex-biases in health and disease. Gene expression from the X chromosome is partly equalized between XX females and XY males by a dosage compensation mechanism called X chromosome inactivation (XCI), which epigenetically silences one of the two X chromosomes in females. However, this inactivation is not absolute – about 12% of X chromosome genes are consistently expressed from the otherwise “inactive” X chromosome (i.e., they “escape” inactivation on X_i_), and another 15% show variable XCI escape across individuals and tissues [15,16].

Escape from XCI is well established for genes located in the Xp and Yp pseudoautosomal regions (PAR1) of the X and Y chromosomes, respectively. These genes tend to be expressed from both sex chromosomes in both sexes (i.e., they are expressed from both the X and Y chromosomes in males; and they escape XCI and are expressed from both X chromosomes in females). Contrary to expectations of equal dosage between males and females, however, PAR1 genes actually tend to be more highly expressed in males [9,16,17]. Although the mechanisms of male-biased PAR1 gene expression remain unclear, this could reflect PAR1 gene expression upregulation in males and/or expression suppression in females via the spreading of XCI [16,18–21]. The latter is supported by observations of lower PAR1 gene expression from the inactive X chromosome (X_i_) than from the active X chromosome (X_a_) in females [16,18–21]. Furthermore, some genes in the PAR2 (e.g., *SPRY3* and *VAMP7*) may be epigenetically silenced on the X_i_ in females and the Y in males [21,22], highlighting the potential for PAR genes to be subject to XCI. **Nevertheless, it remains unclear how the magnitude of inactivation varies across PAR1 (hereafter referred to as “PAR”) genes and tissues and how these patterns may shape sex differences in PAR gene expression.**

Non-PAR X-linked (NPX) genes that escape XCI often show female-biased expression [16,23], but the latter is only an indirect proxy for escape that can lead to false positives [16,24]. Female-biased expression can also arise from mechanisms independent of XCI, such as upregulation from X_a_ in females greater than that from the one X in males (X_M_), similar to mechanisms driving female-biased expression on autosomes [16,24]. Furthermore, escape from XCI may not necessarily result in female-biased expression if, for example, upregulation from the X_M_ in males is even greater than the combined expression from X_a_ and X_i_ females [24]. The most direct evidence of XCI escape is biallelic expression – expression from both X_a_ and X_i_ in females [24].

Studies using allelic expression (AE) have confirmed that escapees tend to exhibit female-biased expression [16] and suggest that higher expression from X_i_ may lead to stronger female-biased expression [16,25,26]. **However, we lack a detailed survey of how variation in the magnitude of escape from XCI relates to the degree of sex-biased expression across NPX genes and tissues throughout the human body.**

Here, I integrated data on allele-specific expression [27], sex-biased gene expression [28], and sex-biased regulatory element activity [29] from across the human body — spanning 25 tissues in the GTEx — to uncover how variation in the strength of XCI influences sex differences in expression. I show that XCI extends beyond the PAR-NPX boundary, and that PAR genes within the same topologically associating domains (TADs) exhibit similar XCI levels, with a cytokine receptor cluster showing especially high escape [30]. These findings were confirmed in analyses of multiple single cell datasets. I also find that the magnitude of inactivation – measured in only three females with skewed XCI [27] – strongly predicts male-biased expression of PAR genes in the GTEx, and that male-biased PAR genes are near regions showing male-biased enhancer activity. Furthermore, I confirm that while many female-biased NPX genes do not escape from XCI, the magnitude of escape predicts female-biased expression for NPX escape genes in the GTEx. The latter are also organized by TADs and near areas of female-biased enhancer activity. This work builds upon previous studies linking sex chromosome dynamics and sex-biased expression by: i) highlighting previously undescribed variation across tissues and genes; and ii) demonstrating that variation in the magnitude of escape from XCI predicts sex-biased expression across different compartments of the sex chromosomes, but only for sex-biased escape genes.

## Methods

### Data collection

#### Allele-specific expression (escape from XCI)

I obtained tissue-specific X-inactivation statuses of X chromosome genes in 30 tissues from a recent analysis of three female GTEx donors with non-mosaic (i.e., highly skewed) X chromosome inactivation [27] (Table S1). A brief summary of the methods is provided here. To identify females with non-mosaic X chromosome inactivation (nmXCI), researchers analyzed heterozygous SNPs on the X chromosome using whole exome sequencing (WES) and RNA-seq data from GTEx. They quantified allelic expression (AE) for each sample, which measures the relative expression from maternal versus paternal alleles at heterozygous sites (AE = |0.5 - (# reference reads / # total reads)|; AE = 0 = half of the reads come from the reference allele and half come from the alternative allele = biallelic expression; AE = 0.5 = all reads come from reference or alternative allele = monoallelic expression). Females with complete, or nearly complete, inactivation of one X chromosome results in highly skewed expression toward one allele (high AE). For each female, they calculated the median AE across all X chromosome genes, excluding the pseudoautosomal region (PAR) and genes previously reported to exhibit variable escape from XCI. Individuals with highly skewed AE (AE > 0.475) were classified as nmXCI, indicating expression from a single parental X. A previously characterized nmXCI individual (UPIC) [16] was included as a positive control. UPIC has extremely high AE values, which are thought to reflect complete skewing towards one X chromosome (i.e., primary skewing). The two females identified in this analysis have slightly greater variation in AE across tissues than UPIC, suggesting clonal selection driven by constitutive genetic variants (i.e., secondary skewing). For each of these samples and in each tissue, each gene in each tissue was categorized as monoallelic (AE > 0.4) or biallelic (AE ≤ 0.4), incorporating the potential consequences of high and low read counts. Throughout, I visualize allelic expression as 1-AE, such that values <0.6 indicate monoallelic expression (silencing by XCI) and values ≥0.6 indicate biallelic expression (i.e., escape from XCI).

Allele-specific expression data was also collected from multiple single cell datasets, including: (i) Garieri and colleagues [31]: X_i_/total across 5 individuals; Tomofuji and colleagues [32]: X_i_/total across multiple datasets (AIDA, Japanese, Tabula Sapiens, XXY, multiome) averaged across samples per tissue or cell type; Tukiainen and colleagues [16]: X_a_/total across 4 individuals; San Roman and colleagues [33]: X_i_/X_a_ across multiple LCL and fibroblast lines (measures adjusted for skewing were used). All values were converted to X_i_/total expression. No PAR genes were included in the data from Garieri and colleagues [31] or from some datasets analysed by Tomofuji and colleagues [32]. Analyses were performed on data averaged across samples and on sample-level data.

#### Sex-biased gene expression

I obtained mean sex differences in expression (i.e., sex-biased expression) for all expressed X chromosome and PAR genes in each of 40 tissues (excluding sex-specific tissues) from a recent analysis of sex chromosome gene co-expression in the GTEx [28] (Table S2). A brief summary of the methods is provided here. Within each gene, expression was modeled as a function of sex, age, and technical variables using limma-voom. To increase power and improve the accuracy of sex effect estimates, multivariate adaptive shrinkage (MASH) was applied [34]. This statistical method combines information across genes and tissues while accounting for uncertainty and correlation in the data. Genes with significant sex-bias (LFSR<0.05) and positive sex effects (β_mash_>0) were categorized as male-biased, while those with significant sex-bias (LFSR<0.05) and negative sex effects (β_mash_<0) were categorized as female-biased. Sex effects on expression reported in this study were very similar to those reported by another large analysis of sex-biased expression in the GTEx [9] (Table S3; Figure S1), although the latter only reported the top 500 sex-biased genes per tissue.

#### Enhancer activity

I obtained sex-biased active regulatory elements (AREs) from a recent analysis of four tissues (brain, heart, muscle, lung; N=387 samples total) from the GTEx data [29] (Table S4). A brief summary of the methods is provided here. The authors performed H3K27ac ChIP–seq on 387 samples from brain, heart, muscle, and lung across 256 GTEx participants. They detected >200K active regulatory elements and used epigenomic coactivity patterns to classify them into 14 regulatory groups. Sex-biased ARE signals were identified by modeling ARE activity as a function of age, tissue-archetype fractions, and technical and clinical factors using limma-voom. AREs with p_adj_<0.2 were considered significantly sex-biased (as in the original publication). Homer was used to annotate sex-biased AREs and identify the gene closest to each.

#### PAR gene data

PAR gene locations were obtained using the R package *biomaRt* [35] (*getBM* function; attributes = start_position and end_position).

#### TAD coordinates

TAD coordinates were obtained from the 3D Genome Browser [36] (for hg38) (“TAD set 1” = dataset A172 from [37]; “TAD set 2” = dataset CD8-positive from [38]).

### Statistical analyses

To test whether allelic expression levels were more similar within TADs than between TADs, I employed a pairwise distance approach. Specifically, for each pair of genes with allele-specific expression data, I calculated their distance as the absolute difference in their AE values (|AE gene_1_ - AE gene_2_|). Gene pairs were then classified based on their genomic organization: pairs where both genes resided in the same TAD were labeled “within-TAD,” while pairs where genes resided in different TADs were labeled “between-TAD.” Comparisons involving the same gene were excluded. I then computed the distribution of pairwise distances for within-TAD versus between-TAD gene pairs. To assess statistical significance, I performed a permutation test in which TAD assignments were randomly shuffled across genes (1,000 iterations), and the difference in median pairwise distances (within-TAD minus between-TAD) was recalculated for each permutation. The observed difference was compared to this null distribution to generate an empirical p-value. These analyses were run among PAR genes only (using GTEx data and both TAD sets 1 and 2; using single cell data and TAD set 1), across NPX genes only (using GTEx data and TAD set 1), and across all X chromosome genes (using GTEx data and TAD set 1).

For the analysis of the whole X chromosome, I also tested whether adjacent TADs exhibited more similar AE values than non-adjacent TADs. I first ordered TADs by their genomic start position and assigned each a sequential rank. For every pair of TADs, I calculated the pairwise absolute differences in allelic expression between genes. Each TAD pair was then annotated by their rank distance (adjacent or non-adjacent TADs). I then summarized the distribution of allelic-expression differences for each category and compared adjacent versus non-adjacent TADs using a Wilcoxon rank-sum test.

I also compared mean escape from XCI (1-AE) between TADs using ANOVA tests (*aov* function in the R package *stats*) and Tukey’s Honest Significant Difference (*TukeyHSD* in the R package *stats*).

I tested for links between gene locations (PAR, NPX), escape status (silenced, escaping), and/or sex-biased expression using binomial tests (*binom.test* function in the R package *stats*; alternative = “greater”). I compared continuous measures of escape from XCI (1-AE) to sex-biased expression (β_mash_) using Spearman’s rank order correlations (*cor.test* function in the R package *stats*). I tested for overlap between sex-biased escape genes and those near regions with sex-biased enhancer activity using Fisher’s exact tests (*fisher.test* function in the R package *stats*).

I also examined relationships between sex-biased expression (β_mash_), XCI escape (1-AE), and mean expression level (normalized gene expression adjusted for age and technical covariates [28], averaged per gene and tissue across XX individuals) using Spearman’s rank order correlations (*ggpairs* function in the R package *GGally*). Mean expression level (per gene and tissue across XX individuals) was positively correlated with the magnitude of XCI escape (1-AE) for male-biased PAR escape genes and female-biased NPX inactive genes, but not for other gene sets (Figure S2). Mean expression level was negatively correlated with the magnitude of sex-biased expression among male-biased PAR escape genes only (Figure S2). These patterns suggest that higher expression level is associated with greater XCI escape and reduced male-biased PAR escape gene expression.

## Results

### XCI spreads into the PAR in females and constrains male-biased expression of PAR genes

To explore the overlap between categorical measures of XCI escape status and sex bias, I first classified each gene in each tissue by its XCI escape status using allelic expression (AE) data from three females with highly skewed XCI in the GTEx [27]. AE measures the relative expression from the inactive (X_i_) and active (X_a_) X chromosomes, with values ranging from 0 to 0.5 (AE=0 implies equal expression from X_i_ and X_a_; AE=0.5 implies all expression is from X_a_). Each gene in each tissue (and individual) was categorized as silenced (0.5≤1-AE<0.6) or escaping XCI (1-AE≥0.6) [4]. Each gene in each tissue was also assigned a sex-biased expression status based on the direction and significance of the estimated sex effects (β_mash_): female-biased (β_mash_<0 and LFSR<0.05), male-biased (β_mash_>0 and LFSR<0.05), or unbiased (LFSR≥0.05) [28].

I confirmed silencing and lack of sex-biased expression for PAR2 gene *SPRY3* (mean 1-AE=0.508 across 3 observations in 2 nmXCI individuals x 2 tissues; sex-bias LFSR>0.05 in 40 tissues) (AE data were not available for other PAR2 genes) [22] and identified possible tissue-specific instances of PAR gene inactivation (*CD99* in two brain tissues, *CD99P1* in artery and muscle, *DHRSX* in muscle, and *PLCXD1* in artery, brain, and thyroid) (Figures S3A-B; Table S5). I also confirmed that PAR genes almost always escaped XCI in females (N=255/266 gene x tissue pairs showed XCI escape, N=7/266 silenced, N=4/266 variable across individuals, binomial test: 255/266, p<2.2e-16) (Figures 1A, S3A-D; Table S5) and were mostly male-biased in their expression (among XCI escape gene x tissue pairs: 197/255 (77.3%) were male-biased, 56/255 (22%) were unbiased, 2/255 (0.8%) were female-biased, binomial test: 197/255, p<2.2e-16; among silenced gene x tissue pairs: 4/7 (57.1%) were male-biased and 3/7 (42.9%) unbiased, binomial test: 4/7, p = 0.05; among gene x tissue pairs with variable escape: 4/4 (100%) were male-biased, binomial test: 4/4, p = 0.06) (Figures 1A, S3C-D; Table S5). Among male-biased PAR genes, those silenced by XCI in females exhibited stronger male-biased expression (t-test: p<0.05) (Figure 1B). Some gene x tissue combinations showed male-biased expression (LFSR<0.05) even with full XCI escape (i.e., nearly equal expression from X_i_ and X_a_; 1-AE>0.99) (Figure S5C), which may suggest additional mechanisms of PAR gene upregulation in males.

**Figure 1.**
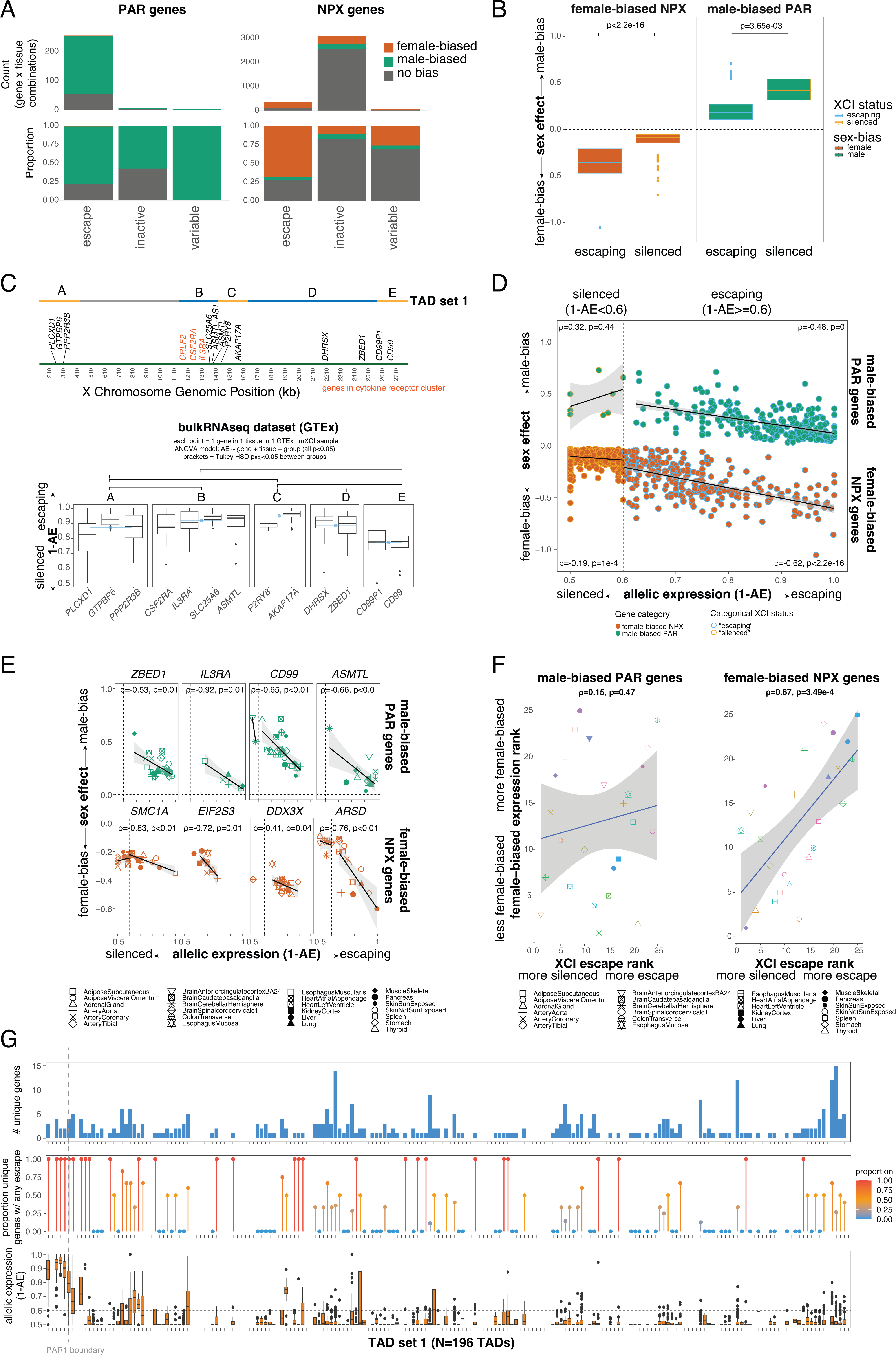
Variation in XCI escape varies across genes and tissues and predicts the magnitude of sex-biased gene expression. A) Left: Bar charts showing the number (top) or proportion (bottom) of PAR gene x tissue observations showing sex-biased expression (see legend) within each tissue-level XCI category (x-axis) (Table S5). Right: Bar charts showing the number (top) or proportion (bottom) of NPX gene x tissue observations showing sex-biased expression (see legend) within each tissue-level XCI category (x-axis) (Table S5). B) Boxplots showing the magnitude of sex-biased expression (β_mash_) for female-biased NPX or male-biased PAR genes that are either silenced by or escape XCI (across gene x tissue observations). C) Top: Positions of PAR genes (using X chromosome annotation) and topologically associated domains (blue and yellow TADs contain PAR genes; grey TAD does not). Bottom: Boxplots of XCI escape (1-AE) for PAR genes, separated by TADs (see top panel). Mean values across all gene x tissue combinations in each TAD are shown in blue. Pairwise significant differences (Tukey HSD p_adj_<0.05) are indicated with brackets. D) Sex-biased expression (β_mash_) (y-axis) versus XCI escape (allelic expression; 1-AE) for female-biased NPX genes (bottom, orange dots) and male-biased PAR genes (top, green dots) (full dataset in Figure S5A). Each dot represents one gene x tissue combination (multiple values were included when AE values from multiple nmXCI individuals were available). Dashed line separates instances of silencing (monoallelic expression, 1-AE<0.6; dots with light orange outline) from XCI escape (biallelic expression, 1-AE≥0.6; dots with light blue outline). Regression lines, confidence intervals, Spearman correlations, and corresponding p-values are shown in each quadrant. E) Similar to Figure 1D, but for individual PAR genes (top row) and NPX genes (bottom row) (see Table S7). Shapes indicate tissue (see legend). F) Left: Tissues ranked by sex bias (median β_mash_) (y-axis) versus XCI (median 1-AE) (x-axis) across tissues (see legend) for male-biased PAR genes. Shapes and colors indicate tissue (see legend). Right: Tissues ranked by sex bias (median β_mash_) (y-axis) versus XCI (median 1-AE) (x-axis) across tissues (see legend) for female-biased NPX genes. Shapes indicate tissue (see legend). G) Top: Bar plot showing the number of unique genes within each TAD (set 1) across the X chromosome. Middle: Dot plot showing the proportion of unique genes with at least one observation of escape within each TAD (set 1) across the X chromosome. Red = higher proportion. Blue = lower proportion. Bottom: Boxplots showing allelic expression (1-AE) values across all genes and tissues within each TAD (set 1) across the X chromosome. Horizontal dashed line represents the threshold for escape (1-AE>0.6). The vertical dashed line spanning all plots represents the PAR1 boundary.

I next examined whether the magnitude of XCI (1-AE) varied among PAR genes. Prior studies proposed that male-biased expression of PAR genes may result from XCI spreading into the PAR [16,18–21], although this phenomenon was initially thought to be limited to *CD99* – the gene closest to the boundary between the PAR and NPX regions of the X chromosome [18]. Interestingly, I found that genes with the strongest silencing were located at both ends of the PAR – near the PAR-NPX boundary (*CD99*, *CD99P1*) and at the telomeric end (*PLCXD1*) – whereas genes located in the center of the PAR exhibited higher levels of escape from XCI (Figure 1C). Analyses of single cell allele-specific expression data also supported these findings (see Methods) (Figure S4). Interestingly, the NPX gene lying right outside the PAR1 boundary (*GYG2*) shows higher levels of escape than *CD99* (Table S1). These patterns are consistent with previous reports of partial silencing of *CD99* and *PLCXD1* [18,20,21], but do not support the expectation that genes closest to the PAR-NPX boundary would exhibit the highest levels of inactivation due to the spread of XCI from the NPX [18].

Genes located within the same topologically associated domains (TADs) exhibited more similar levels of XCI escape (1–AE) than genes located in different TADs (TAD set 1: median distance between TADs = 0.088, within TADs = 0.080, p_perm_ = 0; TAD set 2: between TADs = 0.096, within TADs = 0.064, p_perm_ = 0; Figure 1C; Methods). Consistent with this pattern, TAD membership explained more variance in escape magnitude than gene identity or tissue (ANOVA; TAD set 1: F=30.99, p<2e-16; gene: F=6.10, p=3.15e-07; tissue: F=2.57, p=1.30e-04; TAD set 2: F=39.87, p<2e-16; gene: F=5.9, p=1.54e-07; tissue: F=2.57, p=1.30e-04). These findings support prior observations in mice that escape genes are spatially clustered and aligned with TAD structure [39]. Results were similar when allelic expression was estimated from single-cell datasets (TAD set 1; Figures 1C, S4). Notably, one high-escape TAD includes a cytokine receptor cluster (*CRLF2, CSF2RA, IL3RA*) [30] (Figures 1C, S4). Together, these results suggest that resistance to XCI within the central PAR is not random but reflects underlying regulatory architecture. Because evolutionary genome rearrangements frequently disrupt TAD boundaries and generate species-specific enhancer–promoter interactions – particularly among immune-related genes [30] – such restructuring may have facilitated both the spatial clustering of PAR escape genes and their specialization in immune functions.

I next tested whether variation in the magnitude of XCI among PAR genes and tissues was associated with the magnitude of male-biased expression (β_mash_). Among observations of XCI escape (1-AE≥0.6) and male-biased expression (LFSR<0.05 and β_mash_>0), stronger inactivation (lower 1-AE values) was significantly associated with stronger male-biased expression (higher β_mash_ values) (ρ=-0.48, p<2.2e-16) (Figure 1D, top right quadrant; Figure S5A). No such association was observed when genes were silenced (1-AE<0.6; ρ=0.32, p=0.44) (Figure 1D, top left quadrant; Figure S5A). These relationships held within tissues (for male-biased PAR escape genes: r<0 and p<0.05 for 10/22 tissues) (Figure S5B; Table S6) and for some individual PAR genes, including *CD99*, *IL3RA*, *ASMTL*, and *ZBED1* (r<0 and p<0.05 for 4/11 genes) (Figure 1E, top row; Table S7; Figure S5D). Consistent with this, tissues with higher levels of PAR gene escape tended to show lower levels of male-biased PAR gene expression, although this relationship was not significant (for male-biased, escaping PAR genes in each tissue: tissues ranked by sex bias (median β_mash_) vs. median XCI (median 1-AE): ρ=0.15, p=0.47) (Figure 1F). For example, the colon and stomach showed high levels of XCI escape and low levels of male-biased PAR gene expression, while the adrenal gland and heart showed low levels of XCI escape and high levels of male-biased PAR gene expression (Figure 1F). Finally, I tested whether sex-biased enhancer activity (measured using H3K27ac; see Methods) near PAR genes was consistent with male-biased expression of these genes. I found that male-biased PAR escape genes were close to areas showing male-biased, but not female-biased, enhancer activity (Fisher’s exact test: OR=73.42, p=1.816e-15) (Figure S6), consistent with loss of histone acetylation (including H3K27ac) during XCI in females [40].

### Tissue-specific escape from XCI increases female-biased expression of NPX genes

I confirmed that XCI escape was strongly associated with female-biased expression: among XCI escape gene x tissue pairs, 67.6% (240/355) were female-biased, while 3.9% (14/355) were male-biased and 28.5% (101/355) were unbiased; among silenced genes x tissues, 11.0% (340/3097) were female-biased, 6.7% (209/3097) were male-biased, and most (2548/3097, 82.3%) were unbiased; among genes x tissues with variable XCI status, 15/58 (25.9%) were female-biased, 3/58 (5.2%) were male-biased, and most (40/58, 69.0%) were unbiased (Fisher’s exact test [female-biased vs. others, XCI escape vs. others]: OR=16.4, p<2.2e-16) (Figures 1A, S3D-E; Table S5). However, consistent with the limitations of using female-biased expression as a proxy for XCI escape, there was substantial discordance between the two measures. Of 595 NPX gene-tissue combinations showing female-biased expression, 340 (57%) were subject to XCI rather than escaping (Figures 1A, S3D-E, S5A). Conversely, of 355 NPX gene-tissue combinations that escape XCI, 101 (28%) did not exhibit sex-biased expression (Figures 1A, S3D-E, S5A). Similarly, female-biased NPX genes that escape XCI tend to show larger sex differences than female-biased NPX genes subject to XCI (t-test: p<0.05), but this pattern was not universal (Figure 1B).

I then examined how continuous variation in escape from XCI (AE) relates to sex-biased expression (β_mash_) across tissues and female-biased NPX genes. Among observations of XCI escape (1-AE≥0.6) and female-biased expression (LFSR<0.05 and β_mash_<0), stronger escape (higher 1-AE values) was significantly associated with stronger female-biased expression (lower β_mash_ values) (ρ=-0.62, p<2.2e-16) (Figure 1D, bottom right quadrant; Figure S5A). A weaker association was observed when genes were silenced (1-AE<0.6; ρ=-0.19, p=1e-4) (Figure 1D, bottom left quadrant; Figure S5A). These patterns held within tissues (for female-biased NPX escape genes: r<0 and p<0.05 for 12/22 tissues) (Figure S5B; Table S6) and for individual genes (across tissues), including *SMC1A*, *ARSD*, *EIF2S3*, *DDX3X,* and *NLGN4X* (r<0 and p<0.05 for 5/25 genes) (Figure 1E, bottom row; Table S7; Figure 5D). Consistent with this, tissues with higher levels of NPX gene escape showed higher levels of female-biased NPX gene expression (for female-biased, escaping NPX genes in each tissue: tissues ranked by sex bias (median β_mash_) vs. median XCI (median 1-AE): ρ=0.67, p=3.49e-4) (Figure 1F). For example, the liver and pancreas showed high levels of XCI escape and high levels of female-biased NPX gene expression, while the adrenal gland and muscle showed low levels of XCI escape and less female-biased NPX gene expression (Figure 1F). Finally, I found that female-biased NPX escape genes were close to areas showing female-biased, but not male-biased, enhancer activity (Fisher’s exact test: OR=Inf, p=11.846e-06) (Figure S6), consistent with maintenance of histone acetylation (including H3K27ac) among XCI escape genes [40].

Similar to the analysis of PAR genes, NPX genes within the same TADs showed more similar levels of escape to each other than to genes in different TADs (TAD set 1; including all PAR + NPX genes and TADs with >=2 genes: observed median distance between TADs = 0.039, within TADs = 0.020, p_perm_ = 0; including NPX genes only and all TADs: observed median distance between TADs = 0.023, within TADs = 0.019, p_perm_ = 0) (Figure 1G). Similarly, NPX genes in adjacent TADs were also more similar to each other than those in non-adjacent TADs (mean distance between non-adjacent TADs = 0.058, adjacent TADs = 0.046, Wilcoxon test: W = 8.195e11; p < 2.2e-16; Figure 1G) and the distance between two TADs (i.e., the number of TADs between them) predicted more dissimilar levels of escape among the genes contained in those TADs (Spearman’s ρ = 0.645; p < 2.2e-16) (Figure 1G). These findings suggest not only that escape from XCI is organized within TADs, but that genes in nearby TADs may also be affected.

Together, these findings demonstrate that escape from XCI varies across genes and tissues and is a key driver of sex-biased expression for PAR and NPX genes. Furthermore, gene- and tissue-specific variation in XCI – measured from just three females – closely predicts population-level sex differences in gene expression [16]. These results provide novel evidence that tissue-specific XCI escape is a major mechanism underlying variation in the magnitude of sex-biased gene expression throughout the human body.

## Discussion

This study dissects a fundamental mechanism contributing to sex differences in gene expression: the magnitude of X chromosome inactivation (XCI) escape. The results presented here demonstrate that variation in the degree of escape from XCI is a key determinant of whether genes on the X chromosome exhibit sex-biased expression, and the magnitude of such bias. This insight bridges a gap in our understanding of how sex-biased gene expression arises from chromosomal and regulatory architecture.

One of the most striking findings is that XCI is not limited by the classical pseudoautosomal boundary and instead spreads into the PAR, a domain often assumed to escape inactivation entirely. Although previous work suggested that genes closest to the PAR-NPX boundary may be subject to XCI [18], the current study suggests instead that XCI varies systematically among PAR genes and tissues. Genes at the edges of the PAR (e.g., *CD99*, *PLCXD1*) exhibit greater silencing, while those within the central PAR – particularly those within a cytokine receptor gene cluster – tend to escape XCI more fully. Notably, this finding is supported by multiple independent single cell datasets. The patterned nature of this variation suggests an underlying influence of higher-order genome organization. I find that genes located within the same topologically associating domains (TADs) exhibit more similar escape patterns than genes in different TADs, implicating 3D chromatin structure as a determinant of XCI spread. Although TADs have been shown to organize NPX escape genes in mice [39], this has not previously been applied to the human PAR or NPX regions. These results point to a coordinated role for enhancer activity and TAD organization in modulating escape from XCI, where TAD boundaries limit the spread of silencing and facilitate local enhancer-promoter interactions, and sex-biased H3K27ac marks reinforce differential expression outcomes. In the PAR, this regulatory architecture may have evolved to prevent large sex differences in the expression of certain genes (e.g., cytokine receptor genes), while facilitating male-biased expression of other PAR genes.

In fact, these data provide compelling evidence that suppressed escape from XCI in females contributes to male-biased expression of PAR genes. This builds on previous findings that male-biased PAR gene expression is likely to reflect partial silencing [16,18–21] by showing that greater inactivation leads to greater male-bias across genes and tissues. Given that PAR genes are male-biased even when escape from XCI is high in females, PAR genes may be further upregulated in males (relative to females), even if PAR expression from X_M_ and Y are similar to each other within males [16]. These findings are especially noteworthy in light of evolutionary dynamics that are unique to the PAR. The PAR recombines between X and Y chromosomes in males and is thought to be a hotspot for sexually antagonistic genes—genes for which optimal expression levels differ between the sexes. However, if this sexual conflict is not resolved, suppression of recombination between the X and Y chromosomes is favored, thereby reducing the PAR [7]. Male-biased expression in this region may therefore represent an evolutionary resolution to sexual conflict [7], achieved either through reduced expression in females (via XCI) or upregulation in males. By constraining expression in females, XCI may act to suppress deleterious effects of PAR genes while preserving recombination and maintaining beneficial functions in males. Interestingly, I also identify regions of male-biased enhancer activity near male-biased PAR genes, supporting the notion that enhancer-level regulation complements XCI-mediated silencing.

In contrast, although female-biased expression of NPX genes is not always associated with escape from XCI, the magnitude of female-biased expression is strongly predicted by increased escape from XCI. This builds upon previous studies linking the magnitudes of XCI and female-biased expression [16,25,26] by showing that this pattern holds for individual genes that show linked variation in both allelic expression and sex-bias across tissues. Furthermore, while previous studies have qualitatively suggested that escape genes are organized by TADs [39], the current study quantitatively demonstrated that genes within the same or adjacent TADs show more similar escape magnitude than genes in different or non-adjacent TADs, respectively. These findings also align with chromatin-level evidence: female-biased NPX escape genes are enriched near enhancers showing female-biased H3K27ac activity, a histone mark associated with active gene expression. This suggests that the regulatory landscape not only enables but reinforces escape from XCI in a sex-specific manner.

A key strength of this analysis is the integration of allele-specific expression data, sex-biased transcriptomic profiles, and epigenomic enhancer data across multiple tissues. Together, these orthogonal lines of evidence converge to support a model in which tissue-specific and gene-specific escape from XCI is a major driver of sex differences in gene expression. However, several limitations must be acknowledged. Most notably, the primary estimates of XCI escape rely on data from only three females with highly skewed XCI. While such samples are rare, they are essential for directly quantifying X_i_ expression. Despite this limited sample size, I observe largely reproducible patterns of escape across tissues and individuals (although there is some interindividual variation in XCI escape for specific gene x tissue combinations; e.g., *FAM199X* escapes XCI in only one out of two samples within each of three tissues). Furthermore, variation in PAR gene escape was replicated in multiple single cell datasets. Future studies with larger cohorts of individuals exhibiting non-random XCI – and expanded tissue coverage – will be essential to validate and refine these findings. Additionally, while comparisons to mouse models may offer insights [41], key differences in XCI dynamics between species must be carefully considered [42]. Furthermore, tissue and cell-type matched allele-specific expression and TAD datasets may be more appropriate. Additionally, genes with low total expression may appear subject to XCI due to insufficient read depth to detect low-level escape. While I did not apply a total read count threshold (as instances of complete silencing were not always associated with the lowest read counts), this represents a potential source of false negatives that should be considered when interpreting gene-level XCI status.

In summary, these findings offer a detailed survey of how different sets of sex chromosome genes contribute to sex-biased biology. By linking variation in the magnitude of XCI escape to tissue-specific sex differences in gene expression, this work advances our understanding of the molecular basis of sex differences in health and disease. Further exploration of how this regulatory variation interacts with environmental and hormonal factors may yield deeper insights into the etiology of sex-biased disorders and inform strategies for personalized, sex-aware medicine.

## Supplementary Figures

**Figure S1 Sex effects are consistent across datasets**

Sex-biased expression (β_mash_) from DeCasien et al. 2025 (y-axis) versus sex-biased expression (β_mash_) from Oliva et al. 2020 (x-axis) for overlapping genes in each tissue (see legend) (the latter only published the top 500 significant genes per tissue).

**Figure S2 Mean expression level is not consistently linked to XCI escape or sex-bias**

Distributions, scatterplots, and Spearman’s rank order correlation results for pairwise comparisons of allelic expression, sex-biased expression, and mean expression level (see Methods). Colors represent gene sets included in Figure 1D (red = female-biased NPX escape genes; green = female-biased NPX inactive/silenced genes; blue = male-biased PAR escape genes; purple = male-biased PAR inactive/silenced genes). *0.01<p<0.05. **0.001<p<0.01. ***p<0.001.

**Figure S3 Female-biased expression is not a perfect proxy for XCI escape**

A) Bar charts showing the number (top) or proportion (bottom) of gene x tissue observations within different categories of XCI escape (see legend) within each gene-level category (x-axis) (Table S1).

B) Tile plot showing gene x tissue-level data from Figure S3A.

C) Bar charts showing the number (top) or proportion (bottom) of PAR gene x tissue observations showing sex-biased expression (see legend) within each tissue-level XCI category (x-axis) (Table S5). Reproduced in Figure 1A.

D) Tile plot showing gene x tissue-level data from Figures S3C and E.

E) Bar charts showing the number (top) or proportion (bottom) of NPX gene x tissue observations showing sex-biased expression (see legend) within each tissue-level XCI category (x-axis) (Table S5). Reproduced in Figure 1A.

**Figure S4 Single cell datasets recover cross-TAD variation in PAR gene escape from XCI**

A) Boxplots of XCI escape (X_i_ / total expression) for PAR genes from single cell data, separated by TADs (see Figure 1C). Data is from 4 datasets (see Figure S4B) and values were averaged across samples (within cell types and datasets). Mean values across observations within each TAD are shown in red. ANOVA results are shown above the plot. Pairwise significant differences (Tukey HSD p_adj_<0.05) are indicated with brackets.

B) Boxplots of XCI escape (X_i_ / total expression) for PAR genes from 4 single cell datasets, separated by TADs (see Figure 1A). Values were averaged across samples (within cell types and datasets). These data are summarized across datasets in Figure S4A. Mean values across observations within each TAD are shown in red.

C) Boxplots of XCI escape (X_i_ / total expression) for PAR genes from single cell data, separated by TADs (see Figure 1A). Data is from 2 datasets (see Figure S4D) for which data was available per sample. Mean values across observations within each TAD are shown in red. ANOVA results are shown above the plot. Pairwise significant differences (Tukey HSD p_adj_<0.05) are indicated with brackets.

D) Boxplots of XCI escape (X_i_ / total expression) for PAR genes from 2 single cell datasets (per sample data), separated by TADs. These data are summarized across datasets in Figure S4C. Mean values across observations within each TAD are shown in red.

**Figure S5 Relationships between XCI escape and sex-biased expression vary across genes and tissues**

A) Similar to Figure 1D, but for all genes (Table S5).

B) Similar to Figure 1D, but for individual tissues (Table S6).

C) Sex-biased expression (β_mash_) (x-axis) for PAR gene x tissues that show nearly full escape from XCI (i.e., nearly equal expression from X_i_ and X_a_; 1-AE>0.99). Shapes indicate tissue (see legend). Color indicates sex-bias (see legend).

D) Similar to Figure 1D, but for individual genes (Table S7). Shapes indicate tissue (see legend). Partially reproduced in Figure 1E.

**Figure S6 Genes that escape XCI and exhibit sex-biased expression also show sex-biased enhancer activity**

Count of sex-biased peaks (Table S4) (female- or male-biased; bottom) near genes in each sex-biased expression (male- or female-biased; left) category and gene set (NPX vs PAR; top).

## Supplementary Tables

Table S1 Allele-specific expression data (from Gylemo et al. 2025)

Table S2 Sex-biased gene expression data (from DeCasien et al. 2025)

Table S3 Correlations between sex-biased gene expression from DeCasien et al. 2025 and Oliva et al. 2020 (Figure S1)

Table S4 Sex-biased enhancer data (from Hou et al. 2023)

Table S5 Combined data from Tables S1 and S2

Table S6 Correlations between XCI escape (1-AE) and sex-biased expression (β_mash_) within each tissue and gene set (note that genes were included twice if multiple AE values were present) (Figure S5B)

Table S6 Correlations between XCI escape (1-AE) and sex-biased expression (β_mash_) within each gene and gene set (note that tissues were included twice if multiple AE values were present) (Figure S5D)

## Data and code availability

All code and data available at https://github.com/coevolving-unit/xci-sex-bias.

## Acknowledgements

I thank Dr. Amber Trujillo and Dr. Clarissa Rocca for feedback on earlier versions of this manuscript.

